# A human-derived two-antibody cocktail confers prophylactic and therapeutic protection against authentic Mpox virus

**DOI:** 10.64898/2026.08.31.748425

**Authors:** Birte Kalveram, Awadalkareem Adam, Jimmy Kapetshi, Meredith Weglarz, George (Giorgi) Babuadze, Tian Wang, Hugues Fausther-Bovendo

**Affiliations:** Department of Microbiology & Immunology, University of Texas Medical Branch, Galveston, Texas, USA; The Sealy Institute of Drug Discovery, University of Texas Medical Branch, Galveston, Texas, USA; Institute for Human Infections and Immunity, University of Texas Medical Branch, Galveston, Texas, USA; Sealy Institute for Vaccine Sciences, University of Texas Medical Branch, Galveston, Texas, USA; Institut National de Recherche Biomédicale, Kinshasa, Democratic Republic of the Congo; Département de Microbiologie, Université de Kinshasa; Department of Pathology, University of Texas Medical Branch, Galveston, Texas, USA; Sealy Center on Lung Disease, Inflammation and Remodeling, University of Texas Medical Branch, Galveston, TX, USA

**Keywords:** Mpox, monoclonal antibodies, 2-mAb cocktail, CAST-Eij mouse model, therapeutic

## Abstract

With sustained human-to-human transmission worldwide, Mpox virus remains a significant global health burden. However, there are no licensed therapeutics against Mpox, with clinical management limited to supportive care and pain management. Given the virus complex life cycles, effective treatments require the inhibition of both mature intracellular virions (MV) and extracellular virions (EV). Here, we describe the isolation of human monoclonal antibodies (mAbs) from antigen specific memory B cell using flow cytometry-based cell sorting. We also characterize the therapeutic potential of 2-mAb cocktails targeting both MV and EV using an *in vitro* neutralization assay and a mouse challenge model. Several developed human 2-mAb cocktails neutralized authentic Mpox *in vitro*. When administered 24 hours before or after Mpox challenge, the lead 2-mAb cocktail inhibited viral loads in mouse tissues, with the exception of the testes. Overall, our study identifies several human 2-mAb cocktails with therapeutic potential for controlling Mpox disease.

**SIGNIFICANCE:** Mpox virus, which has emerged as a significant public health threat, is now endemic to several countries worldwide. Existing Mpox vaccines offer only partial protection, and the primary antiviral, tecovirimat, has demonstrated no clinical benefits, highlighting an urgent need for novel therapeutics. Recognizing the limitations of animal-derived antibodies for therapeutic use, we isolated human antibodies and demonstrated the therapeutic potential of two-antibody cocktails using an in vitro neutralization assay and a mouse challenge model. Importantly, the two antibodies within our lead cocktail were predicted to bind to conserved epitopes on their Mpox targets. Overall, this study documents the development of a human two-antibody cocktail with both prophylactic and therapeutic potential against Mpox, offering a promising new avenue for clinical intervention.

## INTRODUCTION

The Mpox virus is a reemerging pathogen that has been responsible for two of the last three public health emergencies of international concern (PHEIC). The 2022-2023 outbreak triggered the first Mpox-related PHEIC, during which more than 100,000 cases, caused by Mpox clade IIb virus, were reported across 115 countries (1–4). In August 2024, a second PHEIC was declared due to the rapid surge in Mpox clade Ib cases in the Democratic Republic of Congo and neighboring countries. To date, more than 45,000 cases of clade Ib have been reported (1). While the number of Mpox clade IIb and Ib cases has steadily declined globally, low but sustained human-to-human transmission is occurring worldwide, including in the Americas, Europe, and Africa (1). Currently, Mpox is predominantly transmitted by contact with infectious lesions on mucous membranes, including during intimate contact (3). Mpox infections are characterized by painful rashes, lymphadenopathy, fever, and muscle aches. The need for hospitalization is heterogeneous, ranging between 1-13% of cases (1, 3).

Current Mpox prevention strategies rely on cross-reactive immunity conferred by historical smallpox immunization and on the immunization of at-risk individuals using a third-generation smallpox vaccine. This vaccine is based on a two-dose regimen of modified vaccinia virus Ankara-Bavarian Nordic (MVA-BN) (3, 5, 6). However, these smallpox vaccines confer only partial protection against Mpox disease. Mpox infection has been described in individuals vaccinated against smallpox during childhood (5, 7), and breakthrough Mpox infections have been reported in individuals who received the two required doses of MVA-BN vaccine (5, 8). Vaccine effectiveness is estimated at 76% in healthy individuals and 66% in immunocompromised individuals (5, 9). Finally, Mpox reinfection has been documented, suggesting that Mpox disease does not provide lifelong protective immunity against subsequent Mpox infection (8, 10). Due to limited Mpox vaccine efficacy and a general increase in vaccine hesitancy, therapeutic countermeasures are needed. Currently, there are no FDA-approved treatments against Mpox disease. The treatment of severe Mpox infection relies on a combination of supportive care and pain management. Historically, tecovirimat, an antiviral approved against smallpox, was used to treat severe Mpox infections. However, recent clinical trials (NCT05534984, NCT05559099) observed no clinical benefits in individuals treated with tecovirimat, highlighting the critical need for the development of effective therapeutics against Mpox (11, 12).

The Mpox virus encodes more than 190 proteins. During its relatively complex life cycle, intracellular mature virions (MV) with a single viral membrane and extracellular enveloped virions (EV) which possess a second viral membrane are produced (2, 13, 14). MV and EV encode distinct surface proteins that are potential targets for neutralizing monoclonal antibodies (mAbs). The MV possess more than 20 membrane proteins, while the EV only express 6 membrane antigens [reviewed in (2)]. As inhibition of both EV and MV is required for effective Mpox treatment, mAb cocktails are considered essential. The importance of antibodies for protection against Mpox infection has been demonstrated by multiple studies. In the non-human primate (NHP) model of Mpox infection, B cell depletion before and after immunization abrogated vaccine efficacy. Conversely, passive transfer with purified immunoglobulin from individuals immunized with live vaccinia vaccine protected NHPs from subsequent Mpox challenge (15). Protective antigens on the surface of both EV and MV were previously described, including the M1R, E8L, H3, A29 surface proteins on Mpox MV and the A35, B6R envelope proteins on Mpox EV (16). The protective efficacy of mAbs targeting these antigens was demonstrated in mouse and rabbit models of orthopoxviruses, including ectromelia virus, vaccinia virus, rabbitpox, and Mpox virus (17–25). It is worth noting that most of the above-described mAbs were derived from immunized animals, including mice and NHPs, limiting their therapeutic potential (18, 22–26). Several groups have isolated human Mpox-specific mAbs from convalescent individuals or from previously immunized individuals (16, 17, 27–31). Among these, most groups isolated antibodies against a single Mpox antigen (17, 27, 28, 30), did not report any in vivo protection (29) or only demonstrated in vivo protection using vaccinia virus, a surrogate virus, rather than authentic Mpox virus (16, 17, 27).

The CAST/EiJ mouse model of Mpox infection can recapitulate differences in disease severity between Mpox clades. Furthermore, the route of Mpox infection also impacts viral distribution and disease severity, with intraperitoneal challenge associated with systemic viral replication and more severe disease (32). As a result, the CAST/EiJ mouse model of Mpox infection is often used to assess the protective efficacy of therapeutics (28, 30, 33, 34).

Here, we describe the isolation of Mpox-specific antibodies from healthy human donors. We demonstrate the in vitro neutralization capacity of 2-mAb cocktails designed to target Mpox EV and MV. As a proof of concept, we also demonstrate the prophylactic and therapeutic potential of one of the developed 2-mAb cocktails in CAST/EiJ mice challenged with Mpox clade IIb.

## RESULTS

### Isolation of human Mpox-specific antibodies

To date, there are no licensed therapeutics against Mpox. Here, we sought to develop a human 2-mAb cocktail targeting Mpox EV and MV with therapeutic potential. Smallpox vaccination was discontinued in 1972 (6). We hypothesized that Mpox-specific B cells could still be isolated from older, smallpox-vaccinated individuals. Buffy coats from 10 human donors were purchased from the Gulf Coast Regional Blood Center (Houston, Texas), and peripheral blood mononuclear cells (PBMCs) were isolated. B cells specific to Mpox A35, B6R, E8L, H3, or M1R were isolated by flow cytometry. Notably, in a previous study, most mAbs targeting these antigens were neutralizing *in vitro* (16). Furthermore, A35 and B6R are present on Mpox EV, while E8L, H3, and M1R are present on Mpox MV (2). To that end, all five Mpox antigens were fluorescently labeled. PBMCs were stained with a combination of Mpox antigens, lineage markers, and a viability dye. From 3 independent sorts, 25, 123, and 174 Mpox-specific, IgG-positive B cells were individually sorted and cultured for 2 weeks (**Figure 1A-B, S1**). These individual B cell cultures were screened for IgG production using an in-house serological assay. Among them, 229 out of 322 (71%) B cells secreted IgG antibodies at concentrations above 200 ng/ml. These B cell supernatants were further analyzed for binding to Mpox antigen by ELISA (**Figure 1B)**. Among these, only 7, 3, 1, 3, and 3 supernatants demonstrated significant binding to Mpox B6R, A35, M1R, H3, or E8L proteins, respectively (**Figure 1C**).

**Figure 1.**
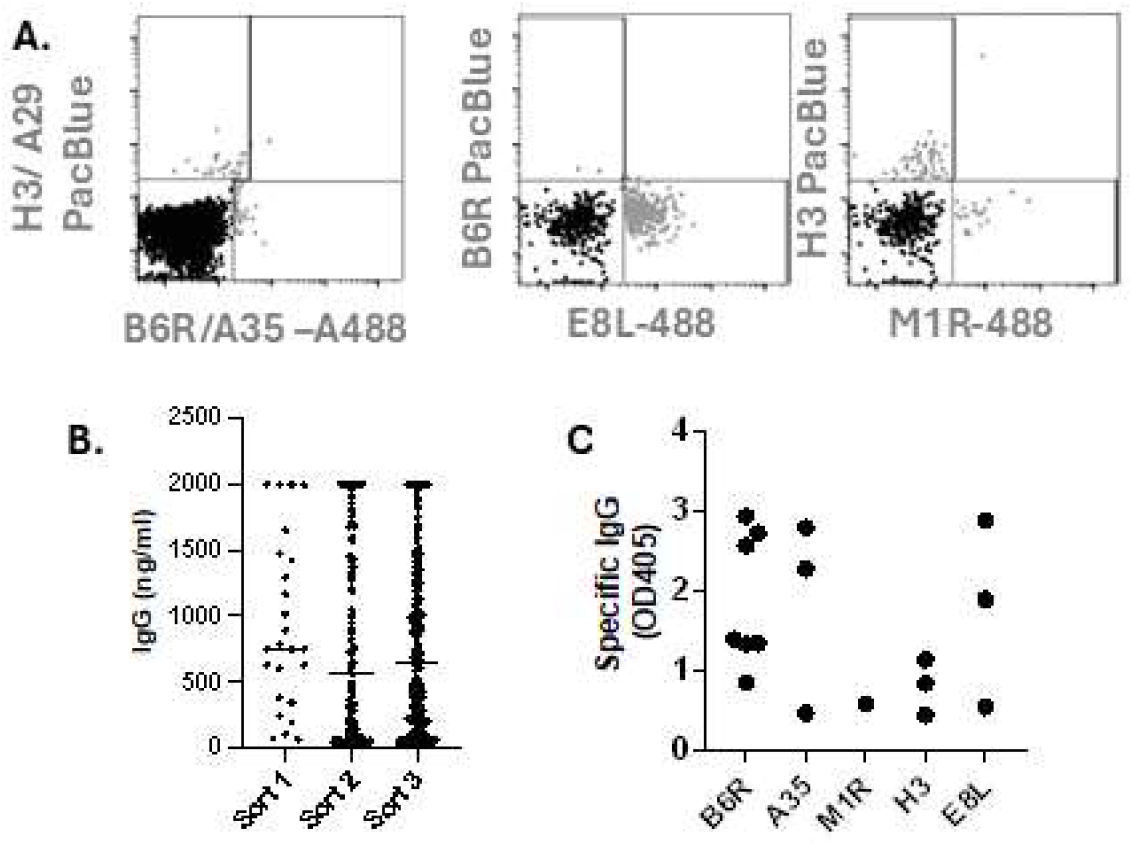
Mpox-specific B cells isolation and culture. Mpox-specific B cells from 10 healthy donors were sorted by flow cytometry (See also Figure S1 for gating strategy). Three independent sorts, including 2, 5, and 5 donors, were performed. Sorted B cells were individually cultured. FACS plots with sorted specific B cells (grey) overlaid onto unstained cells (black) are depicted (**A**.). Using the supernatant from the individual B cell culture, IgG secretion level (**B**.) and binding to various Mpox antigens (**C**.) were tested by ELISA. (**C**.) Only mAbs with binding OD > 0.4 are illustrated. For serological assays, individual dilutions of all samples were run in triplicate.

The variable domains of the 17 antibodies of interest were amplified from the cognate B cell pellets by nested RT-PCR, followed by Sanger sequencing. From these B cells, matching heavy and light chain sequences were generated for 9 unique antibodies. The IGHV4-4*02 and IGKV3-20*01 germlines were disproportionately represented, with 33% and 44% of antibodies derived from these heavy and light chain germlines, respectively (**Figure 2A**). These 9 unique antibodies were cloned, produced by transient transfection of Expi293 cells, and purified using protein A agarose. The binding efficacy of the 9 purified antibodies was determined by ELISA. High binding efficacy was observed against EV antigens, with EC50 values ranging between 8.2 and 11.4 ng/ml for A35-specific antibodies, and between 7.0 and 9.9 ng/ml for B6R-specific antibodies. A larger range of binding efficacy was observed against MV antigens, with binding EC50 values of 8.2, 11.9, and 84 ng/ml against Mpox E8L, H3, and M1R, respectively (**Figure 2B**).

**Figure 2.**
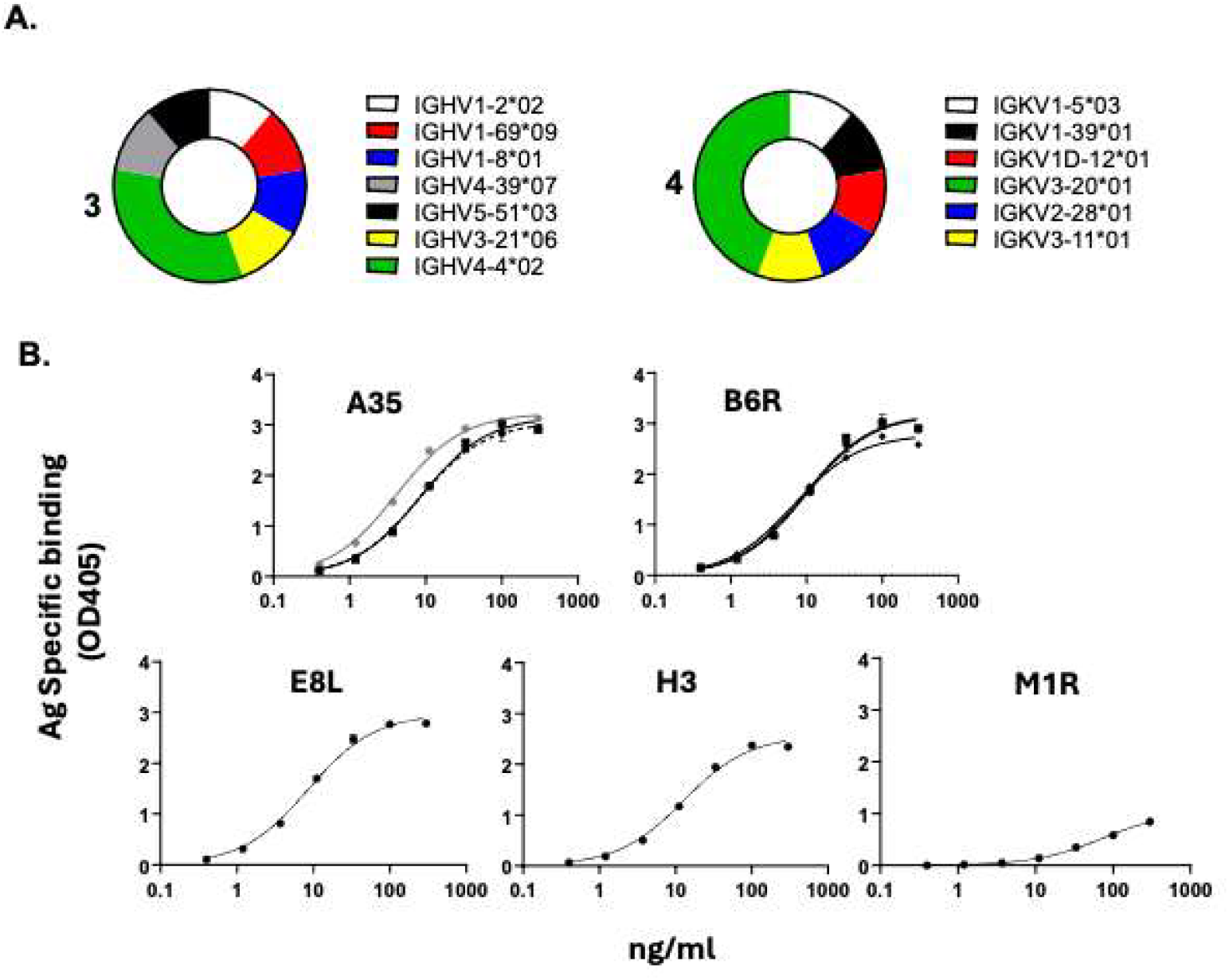
Genetic diversity and binding affinity of isolated Mpox-specific antibodies. The germline usage for the heavy (left) and light (right) chain for the 9 isolated mAbs, determined using IgBlast, are depicted. Germline usage count above 1 is indicated (**A**.). The binding efficiency of antibodies targeting Mpox A35 (n=3), B6R (n=3), E8L (n=1), H3 (n=1), and M1R (n=1) was measured by ELISA. Individual dilutions were evaluated in triplicate (**B**.).

### *In vitro* neutralization capacity of 2-mAb cocktails against Mpox

To assess the functionality of the developed mAbs, their in vitro neutralization capacity was evaluated against authentic clade IIb Mpox virus (USA/MA001/2022). Most antibodies (6 out of 9) targeted Mpox EV proteins (A35 or B6R), while the remaining 3 antibodies targeted Mpox MV antigens. From these 9 antibodies, 18 individual 2-mAb cocktails targeting both MV & EV were derived. The neutralizing capacity of these 18 cocktails was evaluated using a plaque reduction neutralization titer (PRNT) assay. The percentage plaque reduction was evaluated at 10 and 1 µg/ml of each antibody (2 and 20 µg/ml total antibody) in the presence of 10% complement. No reduction in viral plaques was observed at either concentration of the control antibody. In contrast, at the highest tested dose, eight 2-mAb cocktails reduced plaque counts by 50% or more, while at the lowest tested dose, only three 2-mAb cocktails displayed more than 50% reduction in viral plaques. All three of these 2-mAb cocktails (C25-C27) included MP5D9, the E8L-specific antibody, in combination with one of the three B6R-specific antibodies (**Figure 3**). Interestingly, at the 10 µg/ml dose, the highest plaque reduction was observed with a 2-mAb cocktail (C35) including MP10A10, the low-affinity anti-M1R antibody, and MP4F5, a B6R-specific antibody (**Figure 3**). Affinity maturation of the MP10A10 antibody is warranted to further improve the potency of this 2-mAb cocktail

**Figure 3.**
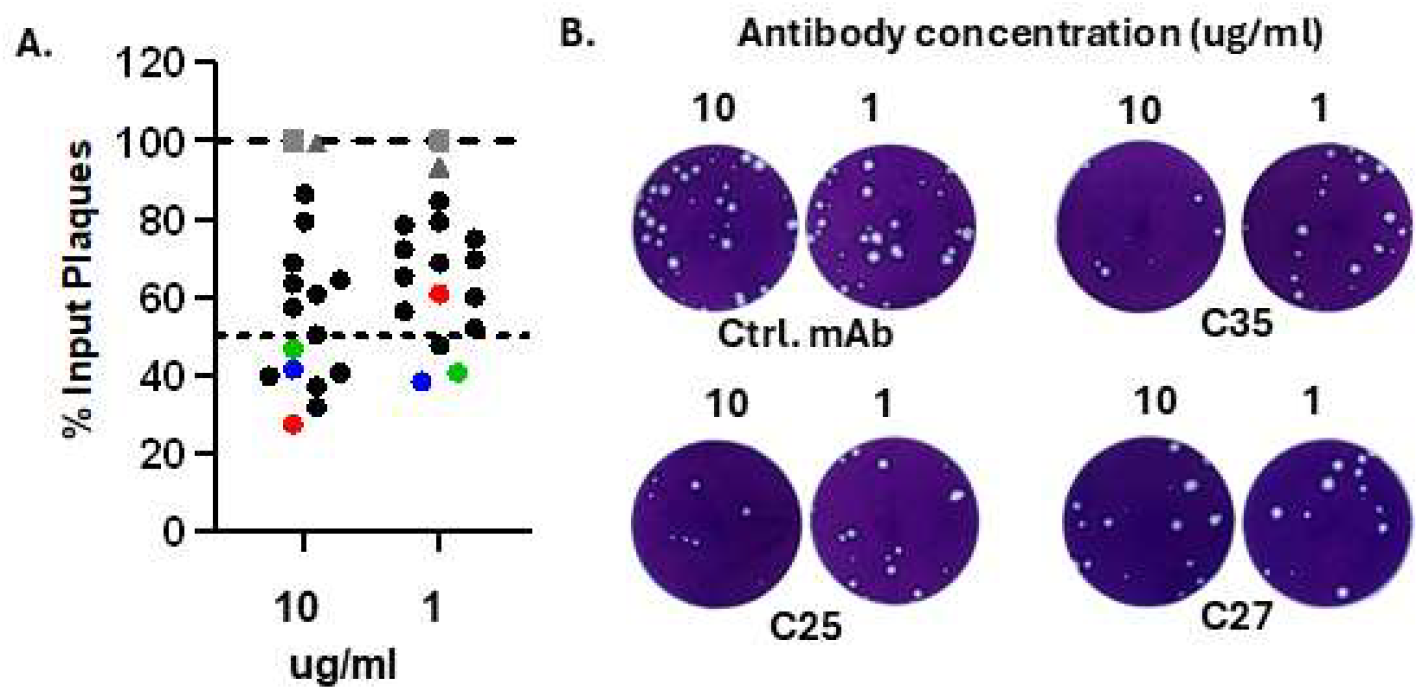
Neutralization capacity of 2-mAb cocktails. Plaque reduction neutralization assay (PRNT) were performed using different 2-mAb cocktail combinations at either 1 or 10 ug/ml of each antibody. The average number of Mpox clade IIb (USA/MA001/2022) viral plaques in the absence of antibody was normalized to 100%. Overall reduction in plaques number of Mpox without antibodies (grey squares), with an irrelevant mAb (grey triangles) or with the various antibody combinations (black circles) are depicted. The C27 (green circle), C25 (blue circle) and C35 cocktails (red circle) were highlighted (**A**.). Representative wells for selected 2-mAb cocktails (denoted C25, C27, and C35) are shown. Each antibody dilution was evaluated in triplicate. (**B**.).

### *In vivo* inhibition of a lead 2-mAb cocktail against Mpox

*In vitro* neutralization does not guarantee *in vivo* efficacy. Thus, inhibition of viral replication by the C27 2-mAb cocktail was evaluated in the CAST/EiJ model of Mpox infection. Among the three most potent 2-mAb cocktails (C25-C27), the C27 cocktail was selected for further evaluation because the two antibodies within this cocktail were produced at high titer, a critical parameter for the manufacture of clinical-grade antibodies (35). The CAST/EiJ model is highly permissive to authentic Mpox virus, with high viral replication observed following intraperitoneal (IP) challenge (32). Male CAST/EiJ mice were infected intraperitoneally with 10^6^ PFU clade IIb Mpox (USA/MA001/2022). Twenty-four hours before or after challenge, animals were treated intraperitoneally with 9 mg/kg of the 2-mAb cocktail (4.5 mg/kg of each antibody). Naïve mice and Mpox-challenged mice treated with 9 mg/kg of an irrelevant mAb were used as controls. Mice were euthanized 6 days after challenge, as this model is not lethal and peak viremia occurred around this timepoint (**Figure 4A, S2**)(32). No significant clinical symptoms were observed in challenged mice, with irrelevant IgG-treated mice showing limited signs of disease (ruffled fur) 6 days after challenge (**Figure 4B**). Accordingly, no weight loss was observed in any of the tested groups (**Figure 4C**). Viral titers were measured by tissue culture infectious dose 50 (TCID50) from tissue homogenates from different groups.

**Figure 4.**
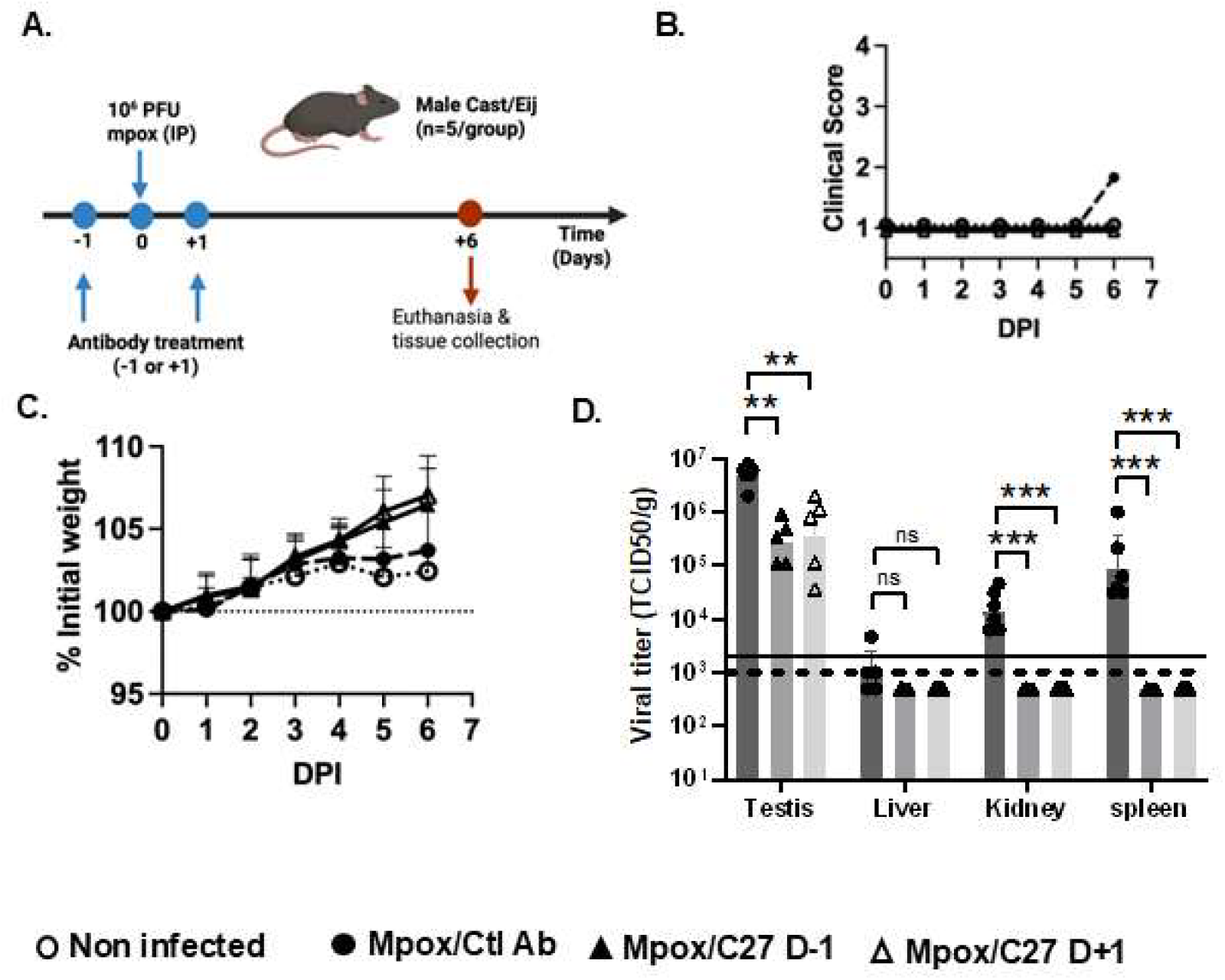
Prophylactic and therapeutic efficacy of the developed 2-mAb cocktail. Seven-week-old male CAST/EiJ mice were challenged intraperitoneally with 10^6^ PFU of clade IIb Mpox (USA/MA001/2022). Animals received, intraperitoneally, 4.5 mg/kg each of MP5D9 and MP12C6 mAb (n=5/group), or 9 mg/kg of control antibody (n=6/group) either 24 hours before or 24 hours after viral challenge. The experimental timeline (**A**.), clinical score (**B**.), and animal weight (mean and standard deviation) (**C**.) were monitored daily for 6 days post infection (DPI). (**D**.) Viral titers were measured 6 days after challenge by TCID50 in the indicated tissues. (**D**.) The mean and standard deviation, * p < 0.05, **p < 0.01, ***p < 0.0001 by One-way ANOVA followed by Dunnett’s test are indicated for each group. (**D**.) The limit of detection, 1000 TCID50/g for liver, kidney, and spleen samples and 2,000 TCID50/g for testis, is indicated by dotted and solid lines, respectively. See also Figure S2.

In the irrelevant mAb group, average viral titers were 5×10^6^, 10^3^, 1.5 ×10^4^ and 9×10^4^ TCID50/g in the testes, liver, kidney and spleen, respectively (**Figure 4D**). In the C27-treated mice, viral titers were below the limit of detection (LOD) in the liver, kidney, and spleen. In kidney and spleen homogenates, the C27 cocktail administered either prophylactically or therapeutically significantly reduced viral titer (p-value < 0.0001). Viral loads in the liver were low, close to or below the limit of detection, across all groups, so no statistically significant difference was observed between the irrelevant mAb and C27-treated groups (**Figure 4D**). In contrast, while the C27 cocktail significantly decreased testicular viral titers (p=0.002 (control vs. day −1 treatment), and p=0.004 (control vs. day +1 treatment)), testicular viral titers remained elevated, with 2.9×10^5^ and 3.7 ×10^5^ TCID50/g average titers in the C27 prophylactic and therapeutic groups, respectively (**Figure 4D**). Overall, the developed 2-mAb cocktail significantly inhibited Mpox viral replication in internal organs but could not clear Mpox from the testes.

### The protective antibodies target conserved epitopes

Finally, the binding epitope of the two antibodies within the C27 cocktail was further evaluated. The structure of each antibody bound to its target was predicted using AlphaFold3 and Protenix-v2 (**Figure 5, S3**). For predictions from both AlphaFold3 and Protenix-v2, the prediction accuracy of the binding interface was high, as illustrated by interface predicted template modeling (ipTM) scores between 0.83 and 0.87 (36, 37). There was a high concordance between the predicted models generated using AlphaFold3 and Protenix-v2, as illustrated by a root mean square deviation (RMSD) values between 2.7 and 4.7 Angstroms for both heavy and light chains (**Figure S3**). Next, the binding footprint of the two antibodies, MP12C6 on B6R and MP5D9 on Mpox E8L, was determined using the PRODIGY server (**Figure 5**). To understand whether the binding epitope of both antibodies was conserved across Mpox clades, all Mpox E8L and B6R sequences available from the National Center for Biotechnology Information (NCBI) were aligned (**supplementary dataset 1 & 2**). The collected Mpox protein sequences included isolates collected between 1979 and 2022, originating from different geographical locations including various states within the US, several African, and European countries. Overall, both Mpox B6R and E8L proteins were highly conserved, with more than 97% homology across the more than 560 aligned sequences. Accordingly, in Mpox E8L, the MP5D9 epitope was conserved, with a single mutation A19T observed near its binding epitope in 32 of the 567 analyzed Mpox isolates (**Figure 6**). Three out of 562 Mpox isolates displayed an S83F mutation within the MP12C6 footprint on the Mpox B6R protein. However, the S83 residue does not appear to be a critical hotspot within the MP12C6 epitope, as 17 amino acids within Mpox B6R display a higher number of interactions with MP12C6. Overall, the binding epitopes of the antibodies within the developed 2-mAb cocktail were highly conserved, and the impact of the observed mutations on the efficacy of the C27 cocktail should be evaluated.

**Figure 5.**
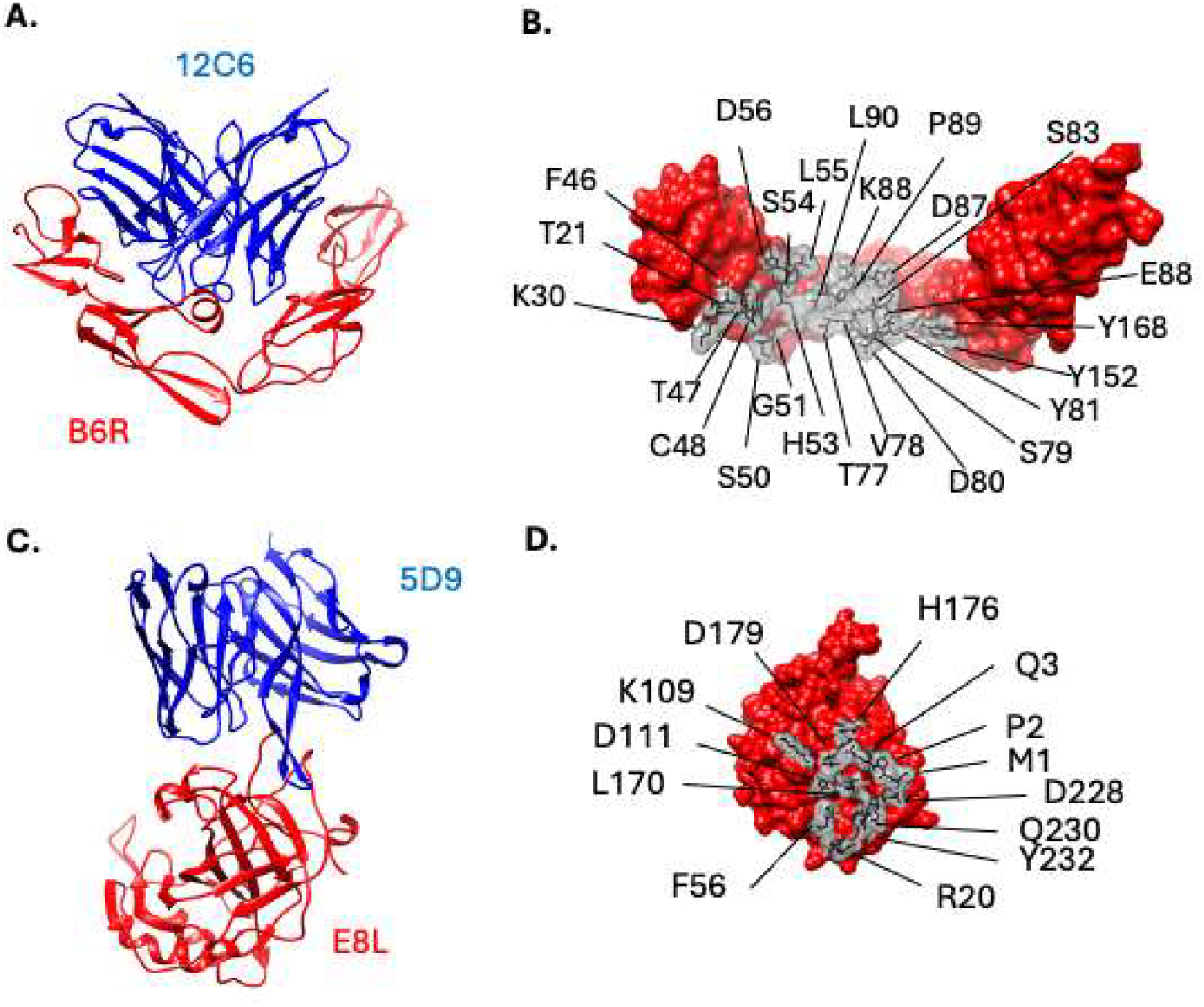
Predicted binding epitope of protective mAbs. The overall AlphaFold3 predicted structure of the MP12C6 (**A**.) and MP5D9 (**C**.) fragment antigen-binding (Fab) region bound to their Mpox target (the B6R and E8L protein, respectively) is depicted. Their epitope footprints, MP12C6 on B6R (**B**.) and MP5D9 on E8L (**D**.), as determined using the PRODIGY server, are shown as sticks. Key amino acids within the Mpox proteins with multiple interactions with MP12C6 and MP5D9 are indicated. Antibodies are depicted in blue while Mpox antigens are red.

**Figure 6.**
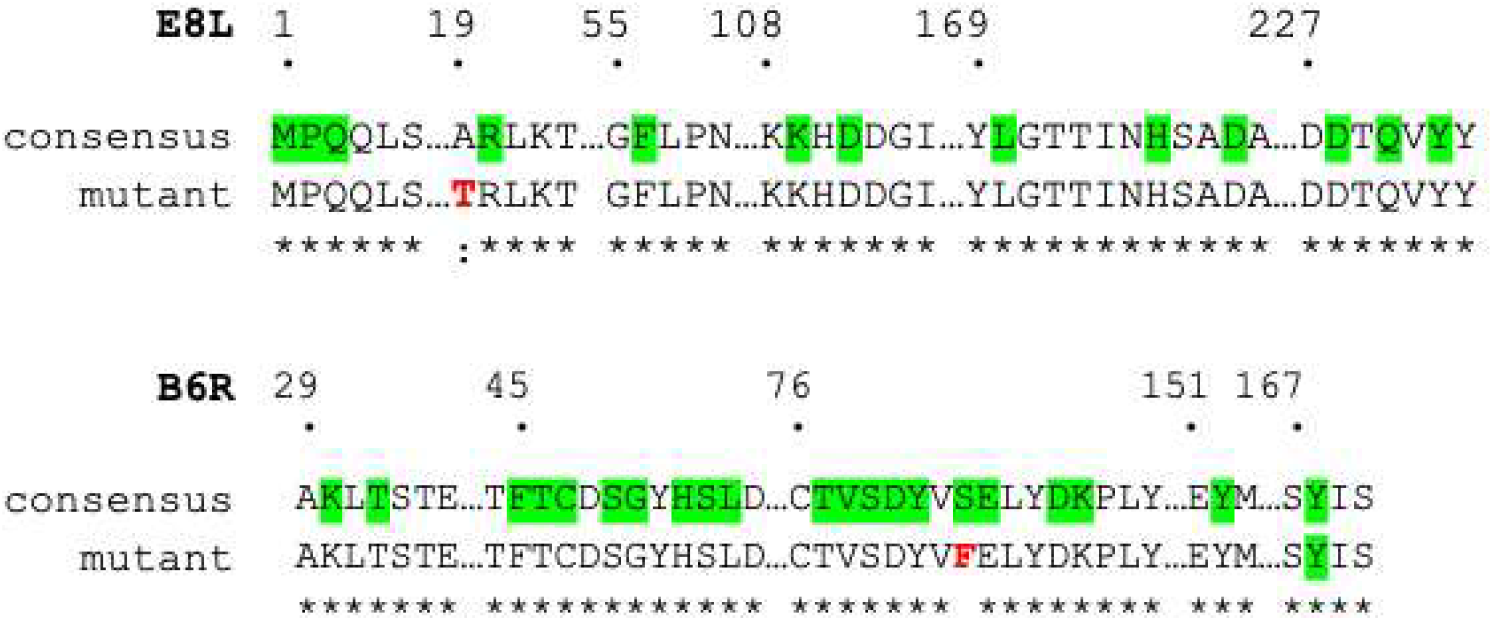
Sequence alignment of the therapeutic mAb epitopes. All available sequences for Mpox E8L (n=567) and B6R (n=562) proteins were obtained from NCBI and aligned using the Clustal Omega multiple sequence alignment server. Observed mutations (red) in or near the MP5D9 and MP12C6 binding hotspots (highlighted in green) are depicted.

## DISCUSSION

The development of therapeutics against Mpox remains a priority for global health. This study documents the isolation of human Mpox-specific antibodies and the development of a protective 2-mAb cocktail. These antibodies were isolated from healthy donors from Texas. However, no clinical information, including the donors’ age or vaccination status, was available. As a result, it is unknown whether the Mpox-specific antibodies originated from previous immunization, including discontinued vaccines; from natural infections; or from cross-reactivity to another pathogen. Prior to this study, a single 2-mAb cocktail of human origin, targeting a different combination of Mpox antigens, namely the B6R and M1R proteins, was reported (31). The protective efficacy of this cocktail was demonstrated in authentic Mpox virus-challenged C57BL/6 mice (31). However, C57BL/6 mice are rarely used to evaluate therapeutics against Mpox. Indeed, the preferred CAST/EiJ mice are more susceptible to authentic Mpox virus than their C57BL/6 counterparts (38). High viral replication and lethality were observed in Mpox clade Ia and IIa-challenged CAST/EiJ mice (32, 38). Given Mpox’s complex life cycles, human antibodies targeting a single Mpox protein are unlikely to provide significant therapeutic benefits in infected individuals (27–30). Due to prohibitive costs associated with Good Manufacturing Practices (GMP) production of mAbs and additional steps required for the licensure of antibody cocktails, larger antibody cocktails are unlikely to be commercially viable (16, 39–41). Similarly, previously isolated, animal-derived, Mpox-specific antibodies will require humanization prior to clinical use (21–25, 33, 42).

The therapeutic efficacy of our developed 2-mAb cocktail was demonstrated in the CAST/EiJ model of infection using authentic Mpox virus administered intraperitoneally. It is worth noting that the therapeutic efficacy of the other developed 2-mAb cocktails was not evaluated in vivo and would require further assessment. The CAST/EiJ model of infection is more stringent than murine models using surrogate viruses such as the vaccinia virus. Indeed, mAb cocktails that reduced viral titer by three logs or more in vaccinia virus-infected mice reduced Mpox viral titer by less than one log in challenged mice (22, 33). In the CAST/EiJ model of Mpox infection, a single dose of the developed 2-mAb cocktail reduced Mpox viral replication when administered prophylactically or therapeutically. To our knowledge, in the CAST/EiJ model of Mpox infection, our 2-mAb cocktail is the only antibody cocktail with demonstrated therapeutic benefits. Using this murine model, reduction in Mpox viral replication was only demonstrated when antibodies were administered prophylactically (28, 33). Of note, Mpox intranasal administration in CAST/EiJ mice was also reported (22, 30, 32, 43). Following intranasal delivery, Mpox predominantly replicates in the airways of challenged mice rather than in internal organs such as the liver and spleen (32). Thus, intranasal Mpox challenge does not recapitulate typical Mpox infection (3).

Compared with 2-mAb cocktails, bispecific antibodies have demonstrated promising therapeutic efficacy in animal models of Mpox, Crimean-Congo hemorrhagic fever (CCHF), and ebolavirus infection (22, 23, 33, 44, 45). Furthermore, compared with a single bispecific antibody, each individual mAb needs to be evaluated before further clinical development of the 2-mAb cocktail (41). Thus, bispecific antibodies derived from the various 2-mAb cocktails will be generated in future studies. Additionally, the in vivo efficacy of the other developed 2-mAb cocktails also deserves further evaluation.

Both Clade Ib and IIb Mpox viruses are currently circulating, so countermeasures should ideally be effective against both viral clades (1). Mpox clade I is considered a select agent in the US, so its use is heavily regulated (https://www.selectagents.gov/sat/list.htm#ftn4). The present study focused on Mpox clade IIb. While this strain was responsible for the 2022-2023 Mpox outbreaks that spread globally, it is a non-select agent (https://www.selectagents.gov/sat/list.htm#ftn4). Whether the developed 2-mAb cocktail is effective against Mpox clade I strains remains to be experimentally confirmed. Of note, both Mpox B6R and E8L are highly conserved, as were the predicted epitopes of both MP12C6 and MP5D9. In more than 560 sequences spanning 40 years, across both Mpox clades and various geographical locations, only a single mutation was observed within or near their binding epitopes. The impact of the S83F mutation within B6R and the A19T mutations within E8L on MP12C6 and MP5D9 binding affinity will require further analysis. However, it is worth noting that the S83F mutation was only detected in 3 isolates (accession number USW05856.1, USW06029.1, USW06202.1) from environmental samples of the residence of a single Mpox-infected United Kingdom resident returning from a trip to Nigeria in 2019, highlighting the scarcity of this mutation (46). The A19T mutation, which is adjacent to the MP5D9 binding epitope, was detected in 5.6% of the analyzed sequences. It is worth noting that this mutation was only detected in samples from Central Africa, with 23 out of 32 (71.9%) of the Mpox isolates with this mutation being collected between 2006 and 2007 and originating from a single human outbreak in the Democratic Republic of Congo (DRC). (47) This mutation was also detected in older DRC Mpox isolates from 1975 (AAY97305.1), 1985 (AKG51453.1), 1996 (Q8V4Y0.1, AAL40563.1, NP_536532.1), 2003 (AAY97104.1), 2008 (AKG51080.1), as well as a 2010 Mpox isolate from the Republic of Congo and a 2016 isolate from Cameroon (WQK80690.1)(48–51). Overall, the protective breath of the generated 2-mAb cocktail will need to be further evaluated.

## Limitations of the study

Mpox can persist in the testes of infected individuals (52, 53). In one study Mpox nucleic acid was present in the semen of over 90% of tested individuals (53). Of additional concern, Mpox was detected in the semen of Mpox-infected individuals after resolution of the skin and mucosal lesions (52). Persistent viruses could contribute to Mpox transmission by unsuspecting male carrier, months after the initial infection. Unfortunately, in the current study and a previous one, protective Mpox-specific antibodies could not eliminate Mpox from the testes of challenged mice (33). Like most therapeutics, antibodies are unable to efficiently cross the blood-testis barrier (54, 55). The development of treatments capable of clearing persistent viruses from the testes of infected individuals will be crucial to reduce Mpox transmission.

In summary, the current study documents the development of a human 2-mAb cocktail with both prophylactic and therapeutic potential against Mpox. Our study warrants further optimization of this 2-mAb cocktail, notably by engineering the isolated antibodies to cross the blood-testis barrier to clear persistent Mpox viruses.

## RESOURCE AVAILABILITY

### Lead contact

Requests for further information and resources should be directed to and will be fulfilled by the lead contact, Hugues Fausther Bovendo.

### Materials availability

All reagents generated in this study are available from the lead contact with a completed materials transfer agreement.

### Data and code availability

Data generated during this study are reported in the manuscript and will be shared by the lead contact upon request.

This paper does not report original code.

Any additional information required to reanalyze the data reported in this paper is available from the lead contact upon request.

## ACKNOWLEDGMENTS

Mpox clade IIb (USA/MA001/2022) was kindly provided by Thomas Ksiazek (The University of Texas Medical Branch). This work was funded by a pilot grant from the Institute for Human Infections and Immunity from the University of Texas Medical Branch, Galveston and by the Sealy Institute of Drug Discovery (to H.F.B.). Molecular representation was performed using UCSF ChimeraX, developed by the Resource for Biocomputing, Visualization, and Informatics at the University of California, San Francisco, with support from National Institutes of Health R01-GM129325 and the Office of Cyber Infrastructure and Computational Biology, National Institute of Allergy and Infectious Diseases.

## AUTHOR CONTRIBUTIONS

Conceptualization, A.A., G.B., and H.F-B.; methodology, B.K. and H.F-B.; Investigation, B.K., A.A., G.B., and H.F-B.; writing – original draft, H.F-B. writing – review & editing, A.A., J.K., G.B., T.W., and H.F-B.; funding acquisition, A.A., T.W. and H.F-B.; resources, J.K.; supervision, T.W. and H.F-B.

## DECLARATION OF GENERATIVE AI AND AI-ASSISTED TECHNOLOGIES IN THE WRITING PROCESS

During the preparation of this work, the authors used Qualified Health AI in order to proofread the manuscript. After using this tool, the authors reviewed and edited the content as needed and take full responsibility for the content of the publication.

## SUPPLEMENTAL INFORMATION

**Document S1. Figures S1–S3**,

**Dataset 1: Alignment of Mpox E8L protein sequences, related to Figure 6 Dataset 2: Alignment of Mpox B6R protein sequences, related to Figure 6**

## EXPERIMENTAL MODEL AND STUDY PARTICIPANT DETAILS

### METHOD DETAILS

#### Human samples

Human blood (Buffy coat) was purchased from the Gulf Coast Regional Blood center (Houston, Texas). Every donor signed an individual inform consent before blood donations. The study was conducted in accordance with the Declaration of Helsinki. No personal information related to the donors was provided to the research team.

#### Animal experiments

Animal experiments were approved by the Animal Care and Use Committee at the University of Texas Medical Branch (protocol # 2410072) and were performed in accordance with the guidelines of the American Association for Laboratory Animal Science.

#### Cell lines

Vero E6 cells (cat# CRL-1586, ATCC, Manassas, VA) were maintained in DMEM (cat# 10013CV, Corning, Corning, NY) supplemented with 5% fetal bovine serum (FBS) (cat# 900-208-500, Gemini Bio, West Sacramento, CA) in a humidified incubator at 37C, 5% CO2. Expi293 cells (Cat# A14527, ThermoFisher scientific, Pittsburgh, PA) were maintained in Expi293™ Expression Medium (Cat# A1435101, ThermoFisher scientific) on an orbital shaker at 125rpm in a humidified incubator at 37C and 7% CO2.

#### Isolation of Mpox-specific mAb

Recombinant Mpox E8L (40890-V08B), A35 (40886-V08H), H3L (40893-V08H1), B6R (40902-V08H), A29 (40891-V08E), M1R (40904-V07H) were all purchased from Sino biological US Inc. (Houston, TX). 100ug of each protein was fluorescently labeled with either Alexa fluor 488 or Pacific Blue using the cognate antibody labeling kit (Cat# A88062, and P30013, both from Thermofisher Scientific). Fluorescently labeled antigens were stored at −80C in single use aliquot.

PBMCs were thawed and stained using 2ug of the fluorescently labeled antigen, a viability marker (Fixable Viability Dye eFluor 780, cat# 65-0865-14, Thermofisher Scientific), and a combination of antibody specific to lineage markers (CD19 (HIB19), CD3 (SP34-2), CD14 (M5E2), IgG (G18-145), IgM (G20-127)), all from BD biosciences. After washing, Mpox-specific B cells were individually sorted into 96 well plates using a FACSARIA Fusion (BD Biosciences). Mpox-specific B cells were sorted and grown for 14 days in RPMI (cat# MT10040CV, Corning) supplemented with 50 ng/ml of recombinant interleukin (IL)-21 (cat# 10584-HNAE, Sino biological US Inc) as previously described (56, 57).

#### Serological assays

Individual wells of half-well, high binding 96 well plates (cat # 07-200-37, Fisher Scientific, Pittsburgh, PA) were coated overnight with 50ng of Mpox proteins (E8L, M1R, B6R, A35 or H3) or unlabeled goat anti-human IgG (cat# NC9830709, Fisher Scientific). Plates were washed then blocked with 5% milk. After 1hr incubation, wells were incubated with the serial dilutions of the samples of interest. After 1hr incubation, plates were washed and incubated with 50ul of 3000-fold diluted HRP conjugated anti-human IgG (cat# A24470, ThermoFisher scientific). After extensive washes, individual wells were incubated with 50µl of ABTS substrate (cat# 50-674-09, Fisher Scientific). Absorbance was read at 405 nm after a final 30mn incubation. To determine IgG concentration, Standard curves generated using human IgG isotype control (cat# 02-7102, ThermoFisher scientific) were used. All samples were run in triplicate.

#### Mpox-specific mAb cloning

Total RNA was extracted from single B cell culture using phenol-chloroform as previously reported (57, 58). The variable heavy and light chain fragments of each antibody were amplified by nested RT-PCR using previously described primers and amplification conditions (59, 60). PCR fragments of heavy and light variable domains were sequenced at the UTMB sequencing core using the Sanger techniques.

#### Antibody production

All antibodies were generated as single-chain variable fragments linked to the fragment crystallizable (Fc) region of a human IgG1 (scFv-IgG) as previously reported (57). Briefly, codon optimized fragment including the heavy chain variable fragment, a [G4S]3 linker, the light chain variable fragment and a IgG1 Fc region were synthesized by GenScript USA inc (Piscataway, NJ). These gene fragments were introduced into pIDV-II, a custom DNA backbone using Gibson assembly (cat# E2611, New England biolabs, Ipswich, MA) (57, 61). The Gibson reactions was used to transform Stellar competent E coli (cat# 636763, Takara bio, San Jose, CA). The generated plasmids were purified and later expanded using the QIAprep Spin Miniprep Kit (cat# 27106) and the QIAfilter Plasmid Midi Kit (cat# 12243), both from Qiagen (Germantown, MD).

Antibodies were produced by transient transfection using the Expi293 expression system kit (Cat# A14635, ThermoFisher scientific). Briefly, 25 ml of Expi293 cell culture were transfected with 25ug of antibody plasmid and 80ul of ExpiFectamine 293 in 2.9ml Optimen (Cat# 31985062, ThermoFisher scientific). A day later, 150ul of transfection enhancer 1 and 1.5ml of transfection enhancer 2 was added to each transfection reaction. Supernatant were collected on day 5. Antibodies (scFv-IgG) were purified using protein A Agarose (cat# 20334, ThermoFisher scientific) as previously reported (59). Antibody concentration was determined by ELISA as described above. Antibody EC50 values were calculated using the AAT Bioquest calculator server (www.aatbio.com/tools/ec50-calculator).

#### Mpox production, titration & Plaque reduction neutralization assay

All experiments involving Mpox virus were performed in the University of Texas Medical Branch (UTMB) Biosafety level (BSL) 3 laboratory. Mpox clade IIb virus (USA/MA001/2022) kindly provided by Dr Thomas Ksiazek and was propagated in Vero E6 cells. Viral preparation generated after a single passage from the stock vial was used for all the experiments described in the present study.

Viral titers were determined by plaque assays. Vero E6 cells were seeded into 12-well plates at 3.5-5 ×10^5^ cells per well the day before infection. 10-fold serial dilutions of the samples were prepared in DMEM supplemented with 2% FBS. Growth medium was removed from Vero E6 monolayers, and each well was inoculated with 100ul of diluted samples. Cells were incubated for 1h with periodical rocking (every 15-20 minutes) during the incubation. After inoculation, each well was overlaid with 1ml of 0.5% Methylcellulose (cat# M352-500, Fisher Scientific) in MEM (cat# 11935046, Fisher Scientific) supplemented with 2% FBS (cat# 900-208-500, Gemini Bio) and 1% Penicillin/Streptomycin (cat# 30001CI, Corning) before returning to the incubator. After 6 days, the overlay was removed and plates were fixed and stained with 0.25% (w/v) Crystal Violet (Thermo Scientific 02286622) in 10% Buffered Formalin (cat# 23245684, Fisher Scientific) for a minimum of 30 minutes before washing and enumerating.

For plaque reduction neutralization titer (PRNT) assay, monoclonal antibodies, approximately 50 PFU of Mpox virus and 10% guinea pig complement (cat# C300-0010, Rockland Immunochemicals, Limerick, PA) were pre-incubated at 37C for 2 hrs. The mAb/Mpox/complement mixture was then added to the Vero E6 cells monolayer for an additional hour. Cells were overlaid with 0.5% methylcellulose for 6 days and stained with crystal violet as described above. All antibodies concentration were evaluated in triplicate.

#### Mpox challenge study

Five- to Six-week-old male CAST/Eij mice (strain #000928) were purchased from Jackson Laboratory (Bar Harbor, ME). Animals were acclimatized for a minimum of 7 days prior to any experimental procedure. For all experiments, animals were challenged intraperitoneally with 106 PFU of Mpox (USA/MA001/2022) in UTMB BSL-3 animal facility. For each experiment, the challenge dose was confirmed by titrating the inoculum. Animals were monitored daily for weight loss and signs of diseases. Investigators were not blinded during animal scoring.

To study Mpox viral kinetic, challenge animals were culled 3 or 6 days post challenge. Naïve mice were used as controls. Following euthanasia, internal tissues including the lung, kidney, spleen, and liver were collected, weighted and homogenized in 1ml of DMEN (cat# 10013CV, Corning) using 5 mm stainless steel beads (cat # 76470-020, SUMMUS industry, Sugar Land, TX) and a TissueLyser (Qiagen). Mpox viral titer was measured by the above-described plaque assay.

To study the efficacy of the developed antibody cocktails, groups of 5 animals were injected intraperitoneally with 4.5mg/kg each of MP5D9 and MP12C6, either 24 hrs before or after Mpox challenge. Groups of 3 animals that received 9 mg/kg of research grade sotrovimab, a SARS-CoV-2 specific mAb (cat# ICH5201, ichor bio, Wantage, United Kingdom) intraperitoneally, either 24 hrs before or after Mpox challenge, were used as controls. Animals were euthanized by exsanguination 6 days after challenge. Internal organs were harvested, weight and homogenized in DMEM as described above. Mpox viral titers were determined using TCID50. Ten-fold serial dilution of tissue homogenate starting at 10mg/ml (spleen, liver and kidney) or 5mg/ml (testes) were added to confluent VEROE6 monolayer for one hour at 37 °C. Tissue homogenates were replaced with DMEM 2% FBS incubate at 37 °C for 1 week. Individual wells were scored for the presence or absence of cytopathic effect (CPE). The Spearman Kärber formula was then used to calculate the TCID50/g (62).

#### Antibody epitope prediction and sequence homology

An overall structure of the Fab of the lead Mpox-specific antibody to their target (Mpox B6R or E8L) was generated using AlphaFold3 using standard parameters (36) and Protenix-v2(37). The latter were executed via the Tamarind Bio platform (https://tamarind.bio). Interacting residues between the antibody and their target Mpox antigen were identified, from the predicted AlphaFold3 structure, using the PRODIGY server (63, 64). Molecular illustration of antibody antigen complex and root mean square deviation (RMSD) calculation between predicted models were performed using the UCSF ChimeraX 1.19 software (65–67).

All E8L and B6R sequences available from NCBI were download. Sequence alignment was performed using the Clustal Omega Multiple sequence alignment (68, 69).

### QUANTIFICATION AND STATISTICAL ANALYSIS

GraphPad Prism 11 was used for statistical analysis. Perform tests; number of replicates are indicated throughout the manuscript both in the figure legends and the results section. p < 0.05 were considered statistically significant. No values were excluded from all the analysis performed.

## REFERENCES

1. World Health Organization, Global Mpox Trends. (2026). Available at: https://worldhealthorg.shinyapps.io/mpx_global/#SEVERITY [Accessed 10 June 2026].

2. H. Yu, W. Resch, B. Moss, Poxvirus structural biology for application to vaccine design. Trends Immunol. 46, 455–470 (2025).

3. O. Mitjà, et al., Monkeypox. Lancet 401, 60–74 (2023).

4. B. Moss, Understanding the biology of monkeypox virus to prevent future outbreaks. Nat. Microbiol. 9, 1408–1416 (2024).

5. A. Hazra, et al., Mpox in people with past infection or a complete vaccination course: a global case series. Lancet Infect. Dis. 24, 57–64 (2024).

6. E. Hammarlund, et al., Multiple diagnostic techniques identify previously vaccinated individuals with protective immunity against monkeypox. Nat. Med. 11, 1005–1011 (2005).

7. K. L. Karem, et al., Monkeypox-induced immunity and failure of childhood smallpox vaccination to provide complete protection. Clin. Vaccine Immunol. 14, 1318–1327 (2007).

8. E. A. G. Faherty, et al., Notes from the Field: Emergence of an Mpox Cluster Primarily Affecting Persons Previously Vaccinated Against Mpox - Chicago, Illinois, March 18-June 12, 2023. Am. J. Transplant 23, 1268–1270 (2023).

9. N. P. Deputy, et al., Vaccine Effectiveness of JYNNEOS against Mpox Disease in the United States. N. Engl. J. Med. 388, 2434–2443 (2023).

10. J. Zeggagh, et al., Second clinical episode of hMPX virus in a man having sex with men. Lancet 401, 1610 (2023).

11. A. R, et al., Tecovirimat for Clade I MPXV Infection in the Democratic Republic of Congo. N. Engl. J. Med. 392, 1484–1496 (2025).

12. J. Zucker, et al., Tecovirimat for the Treatment of Mpox. N. Engl. J. Med. 394, 884–895 (2026).

13. G. L. Smith, A. Vanderplasschen, M. Law, The formation and function of extracellular enveloped vaccinia virus. J. Gen. Virol. 83, 2915–2931 (2002).

14. R. C. Condit, N. Moussatche, P. Traktman, In A Nutshell: Structure and Assembly of the Vaccinia Virion. Adv. Virus Res. 65, 31–124 (2006).

15. Y. Edghill-Smith, et al., Smallpox vaccine-induced antibodies are necessary and sufficient for protection against monkeypox virus. Nat. Med. 11, 740–747 (2005).

16. I. Gilchuk, et al., Cross-Neutralizing and Protective Human Antibody Specificities to Poxvirus Infections. Cell 167, 684–694.e9 (2016).

17. X. Gu, et al., Protective Human Anti-Poxvirus Monoclonal Antibodies Are Generated from Rare Memory B Cells Isolated by Multicolor Antigen Tetramers. Vaccines (Basel). 10 (2022).

18. J. C. Ramírez, E. Tapia, M. Esteban, Administration to mice of a monoclonal antibody that neutralizes the intracellular mature virus form of vaccinia virus limits virus replication efficiently under prophylactic and therapeutic conditions. J. Gen. Virol. 83, 1059–1067 (2002).

19. L. Crickard, et al., Protection of rabbits and immunodeficient mice against lethal poxvirus infections by human monoclonal antibodies. PLoS One 7 (2012).

20. M. H. Matho, et al., Structural and Functional Characterization of Anti-A33 Antibodies Reveal a Potent Cross-Species Orthopoxviruses Neutralizer. PLoS Pathog. 11, 1–25 (2015).

21. M. Li, et al., Three neutralizing mAbs induced by MPXV A29L protein recognizing different epitopes act synergistically against orthopoxvirus. Emerg. Microbes Infect. 12 (2023).

22. M. Li, et al., Bispecific antibodies targeting MPXV A29 and B6 demonstrate efficacy against MPXV infection. J. Virol. 99 (2025).

23. Z. Ren, et al., Identification of mpox M1R and B6R monoclonal and bispecific antibodies that efficiently neutralize authentic mpox virus. Emerg. Microbes Infect. 13 (2024).

24. Z. Chen, et al., Chimpanzee/human mAbs to vaccinia virus B5 protein neutralize vaccinia and smallpox viruses and protect mice against vaccinia virus. Proc. Natl. Acad. Sci. U. S. A. 103, 1882–1887 (2006).

25. E. M. Mucker, et al., Intranasal monkeypox marmoset model: Prophylactic antibody treatment provides benefit against severe monkeypox virus disease. PLoS Negl. Trop. Dis. 12 (2018).

26. E. J. Wolffe, S. Vijaya, B. Moss, A myristylated membrane protein encoded by the vaccinia virus L1R open reading frame is the target of potent neutralizing monoclonal antibodies. Virology 211, 53–63 (1995).

27. R. Zhao, et al., Two noncompeting human neutralizing antibodies targeting MPXV B6 show protective effects against orthopoxvirus infections. Nat. Commun. 15 (2024).

28. B. Ju, et al., Structurally conserved human anti-A35 antibodies protect mice and macaques from mpox virus infection. Cell 188, 6253–6265.e14 (2025).

29. B. Zhou, et al., Two long-lasting human monoclonal antibodies cross-react with monkeypox virus A35 antigen. Cell Discov. 9 (2023).

30. R. F. Fantin, et al., Human monoclonal antibodies targeting A35 protect from death caused by mpox. Cell 188, 6236–6252.e18 (2025).

31. Y. Qu, et al., Generation and characterization of neutralizing antibodies against M1R and B6R proteins of monkeypox virus. Nat. Commun. 16 (2025).

32. J. L. Americo, P. L. Earl, B. Moss, Virulence differences of mpox (monkeypox) virus clades I, IIa, and IIb.1 in a small animal model. Proc. Natl. Acad. Sci. U. S. A. 120 (2023).

33. R. Zhao, et al., Anti-M1R/B6R antibody characterization and bispecific design for enhanced orthopoxvirus protection. EMBO Mol. Med. 17, 2713–2734 (2025).

34. H. Tamir, et al., Synergistic effect of two human-like monoclonal antibodies confers protection against orthopoxvirus infection. Nat. Commun. 15 (2024).

35. Y. Xu, et al., Structure, heterogeneity and developability assessment of therapeutic antibodies. MAbs 11, 239–264 (2019).

36. J. Abramson, et al., Accurate structure prediction of biomolecular interactions with AlphaFold 3. Nature 630, 493–500 (2024).

37. Y. Zhang, et al., Protenix-v2: Broadening the Reach of Structure Prediction and Biomolecular Design. bioRxiv 2026.04.10.717613 (2026). 10.64898/2026.04.10.717613.

38. J. L. Americo, B. Moss, P. L. Earl, Identification of wild-derived inbred mouse strains highly susceptible to monkeypox virus infection for use as small animal models. J. Virol. 84, 8172–8180 (2010).

39. K. J. Whaley, L. Zeitlin, Emerging antibody-based products for infectious diseases: Planning for metric ton manufacturing. Hum. Vaccin. Immunother. 18 (2022).

40. H. Fausther-Bovendo, G. Kobinger, The road to effective and accessible antibody therapies against Ebola virus. Curr. Opin. Virol. 54 (2022).

41. D. Krieg, G. Winter, H. L. Svilenov, It is Never Too Late for a Cocktail - Development and Analytical Characterization of Fixed-dose Antibody Combinations. J. Pharm. Sci. 111, 2149–2157 (2022).

42. W. L. Ling, W. H. Lua, S. Ken-En Gan, Sagacity in antibody humanization for therapeutics, diagnostics and research purposes: considerations of antibody elements and their roles. Antib. Ther. 3, 71–79 (2020).

43. D. Akazawa, et al., Bivalent single-domain antibodies show potent mpox virus neutralization through M1R antigen. Commun. Biol. 8 (2025).

44. J. M. Fels, et al., Protective neutralizing antibodies from human survivors of Crimean-Congo hemorrhagic fever. Cell 184, 3486–3501.e21 (2021).

45. A. Z. Wec, et al., A “Trojan horse” bispecific-antibody strategy for broad protection against ebolaviruses. Science 354, 350–354 (2016).

46. B. Atkinson, et al., Infection-competent monkeypox virus contamination identified in domestic settings following an imported case of monkeypox into the UK. Environ. Microbiol. 24, 4561–4569 (2022).

47. J. R. Kugelman, et al., Genomic variability of monkeypox virus among humans, Democratic Republic of the Congo. Emerg. Infect. Dis. 20, 232–239 (2014).

48. A. M. Likos, et al., A tale of two clades: monkeypox viruses. J. Gen. Virol. 86, 2661–2672 (2005).

49. Y. Nakazawa, et al., A phylogeographic investigation of African monkeypox. Viruses 7, 2168–2184 (2015).

50. S. N. Shchelkunov, et al., Human monkeypox and smallpox viruses: genomic comparison. FEBS Lett. 509, 66–70 (2001).

51. M. G. Reynolds, et al., Detection of human monkeypox in the Republic of the Congo following intensive community education. Am. J. Trop. Med. Hyg. 88, 982–985 (2013).

52. A. Antinori, et al., Epidemiological, clinical and virological characteristics of four cases of monkeypox support transmission through sexual contact, Italy, May 2022. Euro Surveill. 27 (2022).

53. J. P. Thornhill, et al., Monkeypox Virus Infection in Humans across 16 Countries - April-June 2022. N. Engl. J. Med. 387, 679–691 (2022).

54. F. Wang, J. Zhang, Y. Wang, Y. Chen, D. Han, Viral tropism for the testis and sexual transmission. Front. Immunol. 13 (2022).

55. E. E. Hager-Soto, A. N. Freiberg, S. L. Rossi, Viral Disruption of Blood-Testis Barrier Precedes Testicular Infection. Viruses 17 (2025).

56. K. S. Cox, et al., Rapid isolation of dengue-neutralizing antibodies from single cell-sorted human antigen-specific memory B-cell cultures. MAbs 8, 129–140 (2016).

57. H. Fausther-Bovendo, et al., Rapid In Vivo Screening of Monoclonal Antibody Cocktails Using Hydrodynamic Delivery of DNA-Encoded Modified Antibodies. Biomedicines 13 (2025).

58. L. S. Toni, et al., Optimization of phenol-chloroform RNA extraction. MethodsX 5, 599–608 (2018).

59. K. Smith, et al., Rapid generation of fully human monoclonal antibodies specific to a vaccinating antigen. Nat. Protoc. 4, 372–384 (2009).

60. H. Fausther-Bovendo, et al., A Candidate Therapeutic Monoclonal Antibody Inhibits Both HRSV and HMPV Replication in Mice. Biomedicines 10 (2022).

61. G. G. Babuadze, et al., A novel DNA platform designed for vaccine use with high transgene expression and immunogenicity. Vaccine 39, 7175–7181 (2021).

62. M. A. Ramakrishnan, Determination of 50% endpoint titer using a simple formula. World J. Virol. 5, 85 (2016).

63. A. Vangone, A. Bonvin, PRODIGY: A Contact-based Predictor of Binding Affinity in Protein-protein Complexes. Bio. Protoc. 7 (2017).

64. L. C. Xue, J. P. Rodrigues, P. L. Kastritis, A. M. Bonvin, A. Vangone, PRODIGY: a web server for predicting the binding affinity of protein-protein complexes. Bioinformatics 32, 3676–3678 (2016).

65. E. C. Meng, et al., UCSF ChimeraX: Tools for structure building and analysis. Protein Sci. 32 (2023).

66. E. F. Pettersen, et al., UCSF ChimeraX: Structure visualization for researchers, educators, and developers. Protein Sci. 30, 70–82 (2021).

67. T. D. Goddard, et al., UCSF ChimeraX: Meeting modern challenges in visualization and analysis. Protein Sci. 27, 14–25 (2018).

68. F. Madeira, et al., The EMBL-EBI Job Dispatcher sequence analysis tools framework in 2024. Nucleic Acids Res. 52, W521–W525 (2024).

69. F. Madeira, et al., Using EMBL-EBI Services via Web Interface and Programmatically via Web Services. Curr. Protoc. 4 (2024).

