## Supplemental Fig 1-3 for "A human-derived two-antibody cocktail confers prophylactic and therapeutic protection against authentic Mpox virus"

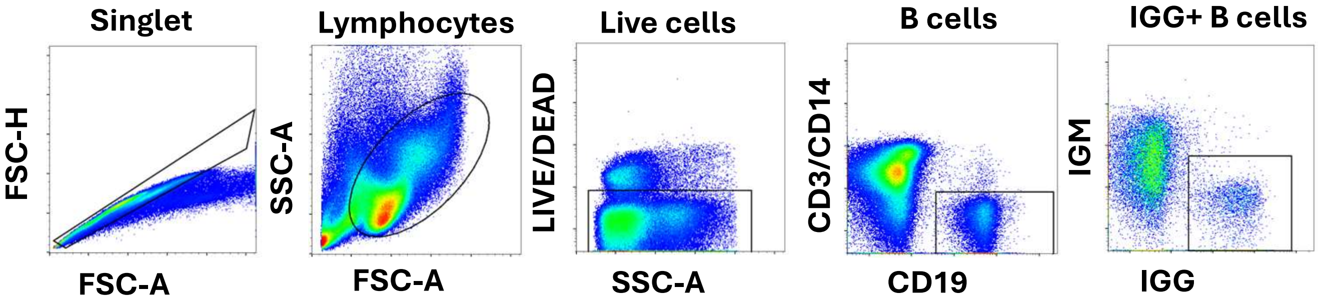


**Figure S1 related to Figure 1: Gating strategy for the isolation of mpox-specific B cells**. IgG+ B Cells were identified based on the following gates: singlet, Lymphocytes, Live (LIVE/DEAD low), B cells (CD19+CD3-CD14-), IgG+ (IgG+, IgM-).

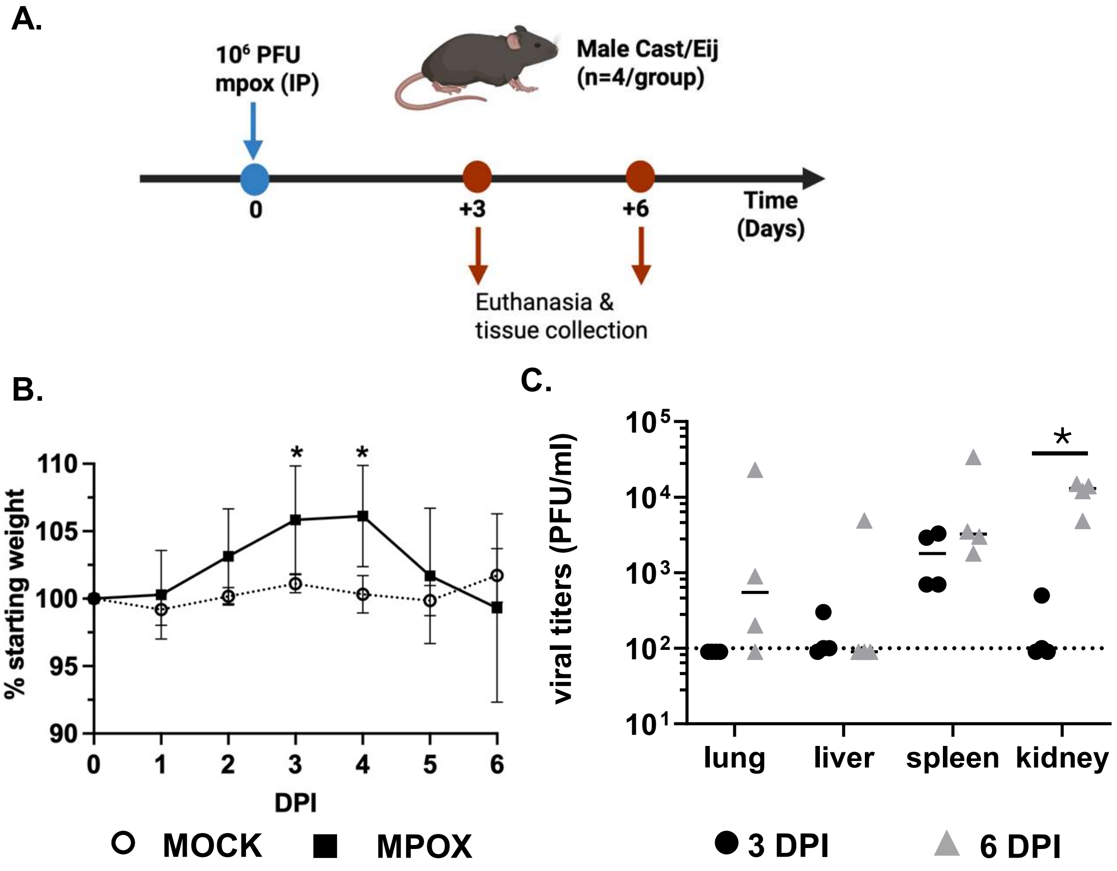


**Figure S2, related to Figure 4: Kinetic of mpox viral replication in Cast/Eij mice. Five** weeks-old male Cast/EiJ mice were infected via the intraperitoneal route with 10^6 PFU of clade IIb Mpox (USA/MA001/2022). Groups of 4 animals were euthanized 3 or 6 days post infection (DPI), and mpox viral titers in the indicated tissues were measured by plaque assay. Each tissue was run in triplicate. The study timeline (A.), animal weight loss (B.) and tissue viral titers (**C**.) are indicated. The mean viral titer, * $p < 0.05$, ** $p < 0.01$ by Welch Student's t test are indicated for each group (**C**.).

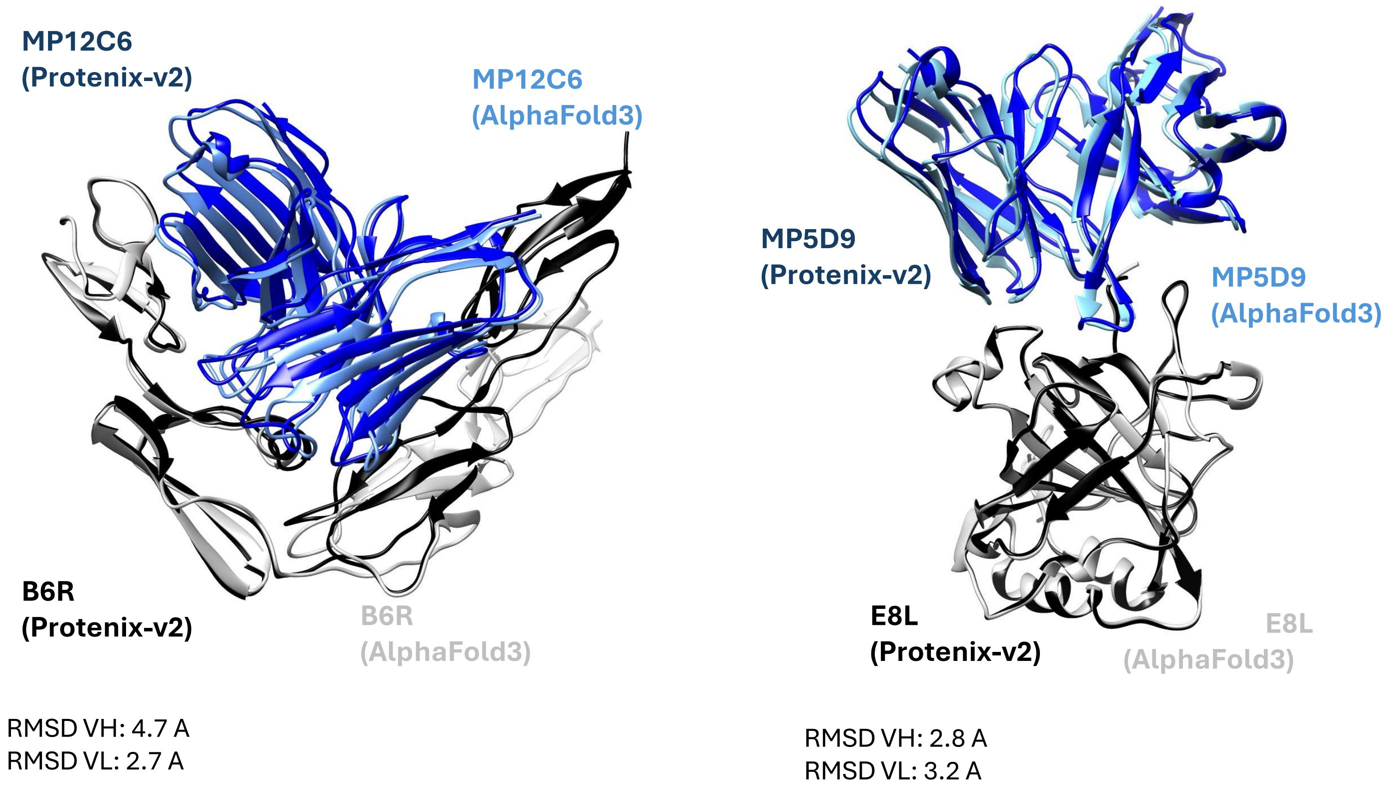


**Figure S3: Comparison of structure prediction models.** The structures of 12C6 bound to Mpox B6R (Left) and 5D9 bound to Mpox E8L (right) were predicted using both AlphaFold3 and Protenix-v2. The Alphafold 3 predicted structures are depicted in lighter colors, while the Protenix-V2 predictions are in darker shades. RMSD value in Angstroms are indicated.
